# Caries Prevalence and Experience in Individuals with Osteogenesis Imperfecta

**DOI:** 10.1101/418806

**Authors:** Mang Shin Ma, Mohammadamin Najirad, Doaa Taqi, Jean-Marc Retrouvey, Faleh Tamimi, Didem Dagdeviren, Francis Glorieux, Brendan Lee, Vernon Sutton, Frank Rauch, Shahrokh Esfandiari

## Abstract

**Objective:** Dentinogenesis Imperfecta (DI) forms a group of dental abnormalities frequently found associated with Osteogenesis Imperfecta (OI), a hereditary disease characterized by bone fragility. The objectives of this study was to quantify the caries experience among different OI-types and quantify how these values change due to DI.

**Methods:** To determine which clinical characteristics were associated with increased CPE in patients with OI, the adjusted DFT scores were used to account for frequent hypodontia, impacted teeth and retained teeth in OI population.

**Results:** The stepwise regression analysis while controlling for all other variables demonstrated the presence of DI (OR 2.43; CI 1.37 to 4.32; p=0.002) as the significant independent predictor of CPE in the final model.

**Conclusion:** This study found no evidence that CPE of OI subjects differs between the types of OI. The presence of DI when controlled for other factors was found to be the significant predictor of CPE.

## Background

Osteogenesis imperfecta (OI) is a genetic connective tissue disorder that is characterized by bone fragility. Although mutations in many genes have been implicated in OI, the large majority of patients have disease-causing variants in COL1A1 or COL1A2, the genes coding for collagen type I ^1,2^. In addition to brittle bones, the clinical manifestations of OI can include developmental disturbances of teeth known as dentinogenesis imperfecta (DI), blue sclera, skin and joint laxity, wormian bones and hearing impairment ^1^.

OI has a wide clinical expression, ranging from very mild to extremely severe with perinatal lethality ^2,3^. This range of severity is mirrored in the traditional Sillence classification into four clinical types ^1,2^. OI type I is the most common and mildest OI type with absence of major bone deformities. OI type II usually results in prenatal or perinatal death. OI type III is the most severe non-lethal form of the disease, and OI type IV has characteristics that are intermediate in disease severity between OI types I and III. Subsequently, this classification has been expanded, first based on distinguishing clinical features (OI types V, VI and VII), and later based on genetic findings ^1,2^.

Teeth affected by DI appear opalescent, with a brown or blue tinge, with early obliteration of pulp chamber and canals, short and slender roots, abnormal dentin, constricted cemento-enamel junction, excessive wear of the teeth and bulbous or tulip shaped crowns ^4,5^.Primary teeth are usually more severely affected than permanent teeth^6^. Tooth enamel can shear off, and the exposed dentin abrades fast. Histologically, the dentin appears amorphous with irregular and obliterated dentinal tubules and a loss of scalloping at the dentino-enamel junction is frequently observed ^5,7^. The frequency of DI varies with the severity of the phenotype and has been reported to affect 31% of individuals with OI type I and up to 86% of patients with OI type III ^5^.

The prevalence of caries in OI has not been assessed in much detail. One study on 40 children with OI types III and IV noted that caries seemed to be a rare finding ^8^. One of the largest study to date assessed caries in 60 children and adolescents with OI types I, III and IV and concluded that the prevalence of caries was similar to that found in the general population ^9^. This is in contrast to the earlier existing hypothesis that altered tooth histology in DI impedes the spread of caries^10^. Furthermore, the OI types represent varying levels of severity of OI with an oro-facial component including an effect in tooth histology, which in turn might have an effect on pattern of dental caries spread. It is presently unknown whether caries in OI varies between OI types and whether it is DI dependent. The hypothesis of this study therefore is that there is no difference in the prevalence of dental caries and experience: between OI population with and without DI; and between different types of OI.

In the present study, we assessed a large group of OI individuals for the presence of caries. Caries is irreversible once the destruction of organic substance has occurred. The lesion can result in a tooth’s exfoliation or extraction, or it can be restored with filling material. These various scenarios are captured by the caries index of decayed, missing and filled teeth (DMFT), which is commonly used of epidemiological studies on caries ^11^. In this study, the relationship between a modified caries index, OI type and presence of DI were studied.

## Methods

The study cohort was recruited through the Brittle Bone Disease Consortium (https://www.rarediseasesnetwork.org/cms/BBD). This consortium is part of a Rare Disease Clinical Research Network that comprises several specialized centers from across North America (Houston, Montreal, Chicago, Baltimore, Portland, Washington DC, New York,Omaha, Los Angeles). One of the projects conducted by the consortium is a natural history study to assess the clinical features of OI. Patients with a diagnosis of OI of any type and any age are eligible to participate. Dental evaluations are offered to participants who are three years of age or older. Patient recruitment started in August 2015. The present evaluation includes data that had been collected on individuals with a clinical diagnosis of OI types I, III and IV until August 2017.

For adults with OI and families with OI affected child, it is often very difficult to access a dental professional for variety of reasons. The level of cooperation among the OI population also varies with level of physical handicap. Many participants travelled from remote places to data collection centers, which are spread across North America. Therefore, to minimize patient discomfort, a decision was made to forego recording of the intra/inter-rater agreement for the baseline year. Caries diagnosis agreements were not assessed in this particular study. To reduce inter-rater variability and for increased sensitivity for caries assessment, the International Caries Detection and Assessment System (ICDAS) ^12^ was employed. The ICDAS scores of 0,1 and 2 were then recoded back to WHO score of ‘0’ for no caries and scores of 3, 4, 5 and 6 to ‘1’ for caries ^13^(figure 1). Any restoration (except sealant) on a tooth was noted as 1 (present). Extensive or multiple restorations on a single tooth was still marked as 1. A tooth showing as both decayed and filled, was still marked as ‘1’. Absence of previous dental records and poor recall of dental history was a recurring theme in individuals with more debilitating forms of OI. This resulted in indetermination of the reason for missing teeth. To take into consideration the prevalence of partial or full hypodontia ^14^ and frequent tooth impactions^15^ in the OI group, that would skew the evaluation of caries experience using DMFT, a mathematical modification was performed to increase the accuracy of the caries experience. This approach was derived from a study of individuals with Down’s Syndrome used by Ulseth et al ^16^ to overcome a similar problem. The DFT (Decayed and filled teeth) score was adjusted by dividing the total number of decayed and filled teeth by the total number for teeth present at the time of the dental examination. The adjusted DFT index is a continuous variable ranging from 0 to 1. Oral hygiene of the participants was evaluated based on the criteria of the debris index of the Oral Hygiene Index ^17^. Mean debris scores of 0.9 and below were classified as good, 1 to 1.9 as fair and score of 2 or greater as poor. The molar classification was also recorded to ascertain whether it has an effect on caries ^18,19^. For the purposes of this study, the dentition of the participants with altered intrinsic color (translucent or opalescent), attrition, altered crown shape and DI typical crown fractures were classified as DI positive.

**Figure 1:**
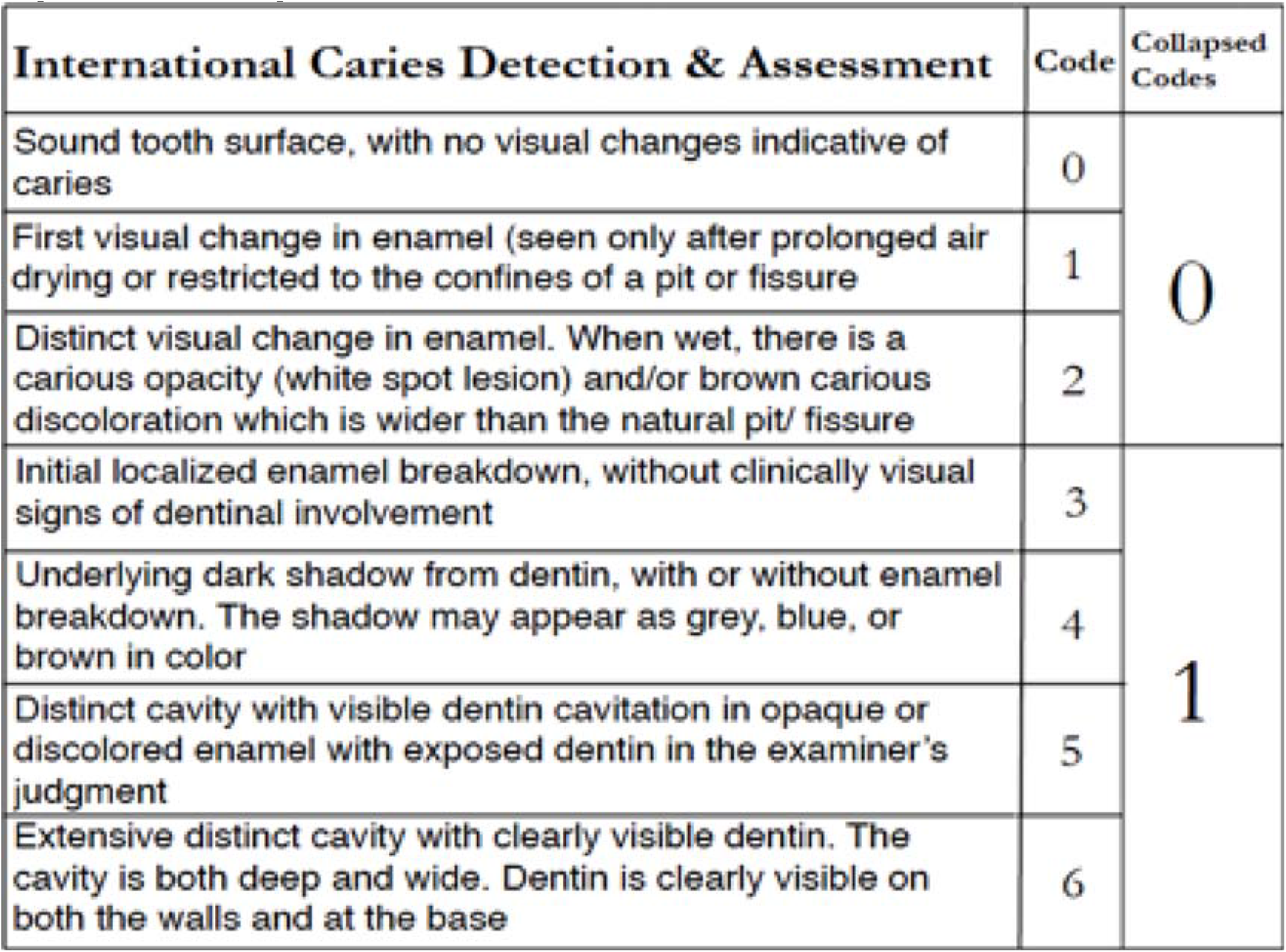
Recoding ICDAS Scores to WHO scores of ‘0’ and ‘1’.

### Data Analysis

Group means were compared by unpaired t tests or ANOVA, as appropriate. The chi square test was used to assess the distribution of categorical data between groups. The relationship between patient characteristics and caries experience was evaluated by logistic regression analysis. Results were expressed as odds ratios (ORs) with 95% confidence intervals (95 % CIs). The effect of potential predictor variable was assessed initially in univariate models followed by stepwise multivariate models. All tests were two-tailed and p-values below 0.05 were considered significant. Stata 13.0 software for Mac (StataCorp.2013. Stata Statistical software; Release 13. College Station, TX: StataCorp LP) was used for statistical analyses.

## Results

A convenience sample of 319 participants was used (median age=18.4years, range=2.8-75.8years) with OI types I (n=150), III (n=68) and IV (n=101). The study population included fewer individuals with mild OI (OI type I, 47%) than participants with moderate to severe OI (OI types III and IV, 53%) (Table 1). As expected, the majority of patients with moderate to severe OI had DI, whereas in OI type I, DI was rare. There was no statistically significant difference in caries prevalence (DT), FT Score and Debris Index between OI types (Table 1). The discrepancies in the sample size of variables between tables 1, 2 and 3 were because of missing data.

**Table 1.** Clinical characteristics of the study population.

| **Independent Variables** | **N** | **All** | **OI Type I** | **OI Type III** | **OI Type IV** |  |
| --- | --- | --- | --- | --- | --- | --- |
| N (m/f) | 319 | 319 (134/185) | 150 (58/92) | 68 (28/40) | 101 (48/53) |  |
| Age (Years); median (range) | 319 | 18.4 (2.8-75.8) | 20.8 (3-75.8) | 14.6 (2.8-57.5) | 17.3 (2.9-71.2) |  |
| DI; n (%) | 303 | 124 (41) | 23 (16) | 51 (82) | 50 (54) | . |
| DT (Score); mean (SD) | 317 | 1.17 (3.14) | 1.31 (3.44) | 1.32 (3.7) | 0.78 (1.9) |  |
| FT (Score); mean (SD) | 316 | 3.98 (5.00) | 4.29 (5.17) | 4.15 (5.41) | 3.55 (4.5) |  |
| Adj. DFT (Score); mean (SD) | 313 | 0.19 (0.22) | 0.19 (0.21) | 0.23 (0.26) | 0.17 (0.19) |  |
| Debris (Score); mean (SD) | 181 | 0.83 (0.63) | 0.74 (0.65) | 0.98 (0.62) | 0.89 (0.58) |  |

P values indicate the significance of the difference between OI types (ANOVA or chi square test).

CPE did not differ between sexes or OI types and was not associated with oral hygiene indices (Table 2). CPE varied with age (with the lowest result in the age group of 8.5 to 13.7 years), and was significantly higher in individuals with DI than in those without DI (Figure 2). To further investigate the clinical characteristics independently associated with CPE, logistic regression analyses were performed (Table 3). Univariate analyses confirmed that the DI was positively associated with CPE, whereas OI type, gender, debris index and molar classification were not significantly related to CPE. These independent variables were entered into a stepwise logistic regression model to ascertain their association with CPE. Presence of DI (OR 2.43; CI 1.37 to 4.32; p=0.002) while controlling for age emerged as significant independent predictors of CPE in our final model.

**Table 2.** Clinical characteristics corresponding to caries prevalence and experience.

|  | **Patients with untreated**<br>**Caries Prevalence (D>0)**<br>**<u>n (%)</u>** | **Patients with**<br>**Caries Experience**<br>**(Adj. DFT>0) <u>n (%)</u>** | **Caries Prevalence**<br>**and Experience**<br>**(Adj. DFT) <u>Mean (SD)</u>** |
| --- | --- | --- | --- |
| Entire group (n=319) | 96 (30) | 223 (70) | 0.19 (0.22) |
| Gender (n = 317) |  |  |  |
| Male | 40 (42) | 89 (40) | 0.18 (0.21) |
| Female | 56 (58) | 134 (60) | 0.20 (0.23) |
| Age (n = 317) |  |  |  |
| 2.8-8.5 Years | 19 (20) | 43 (19) | 0.18* (0.25) |
| 8.5-13.7 Years | 17 (18) | 42 (19) | 0.09* (0.13) |
| 13.7-23.4 Years | 32 (33) | 65 (29) | 0.17* (0.17) |
| 23.4 & Above | 28 (29) | 73 (33) | 0.34* (0.23) |
| OI (n = 317) |  |  |  |
| Type I | 44 (47) | 106 (48) | 0.20 (0.21) |
| Type III | 18 (19) | 48 (22) | 0.23 (0.26) |
| Type IV | 32 (34) | 66 (30) | 0.17 (0.19) |
| DI (n = 305) |  |  |  |
| No | 58 (61) | 116 (54) | 0.16** (0.18) |
| Yes | 37 (39) | 98 (46) | 0.23** (0.25) |
| Debris Index (n = 310) |  |  |  |
| Good | 26# (28) | 80 (64) | 0.18 (0.20) |
| Fair | 23# (25) | 40 (32) | 0.16 (0.17) |
| Poor | 43# (47) | 5 (4) | 0.21 (0.21) |
| Molar Classification (n = 297) |  |  |  |
| I | 23 (2) | 106 (48) | 0.16 (0.21) |
| II | 9 (10) | 48 (22) | 0.20 (0.26) |
| III | 64 (68) | 66 (30) | 0.20 (0.20) |

Statistical Tests: \*p<0.05 by ANOVA; \*\*p<0.05 by t test; #p=0.05 by chi square test

**Figure 2.**
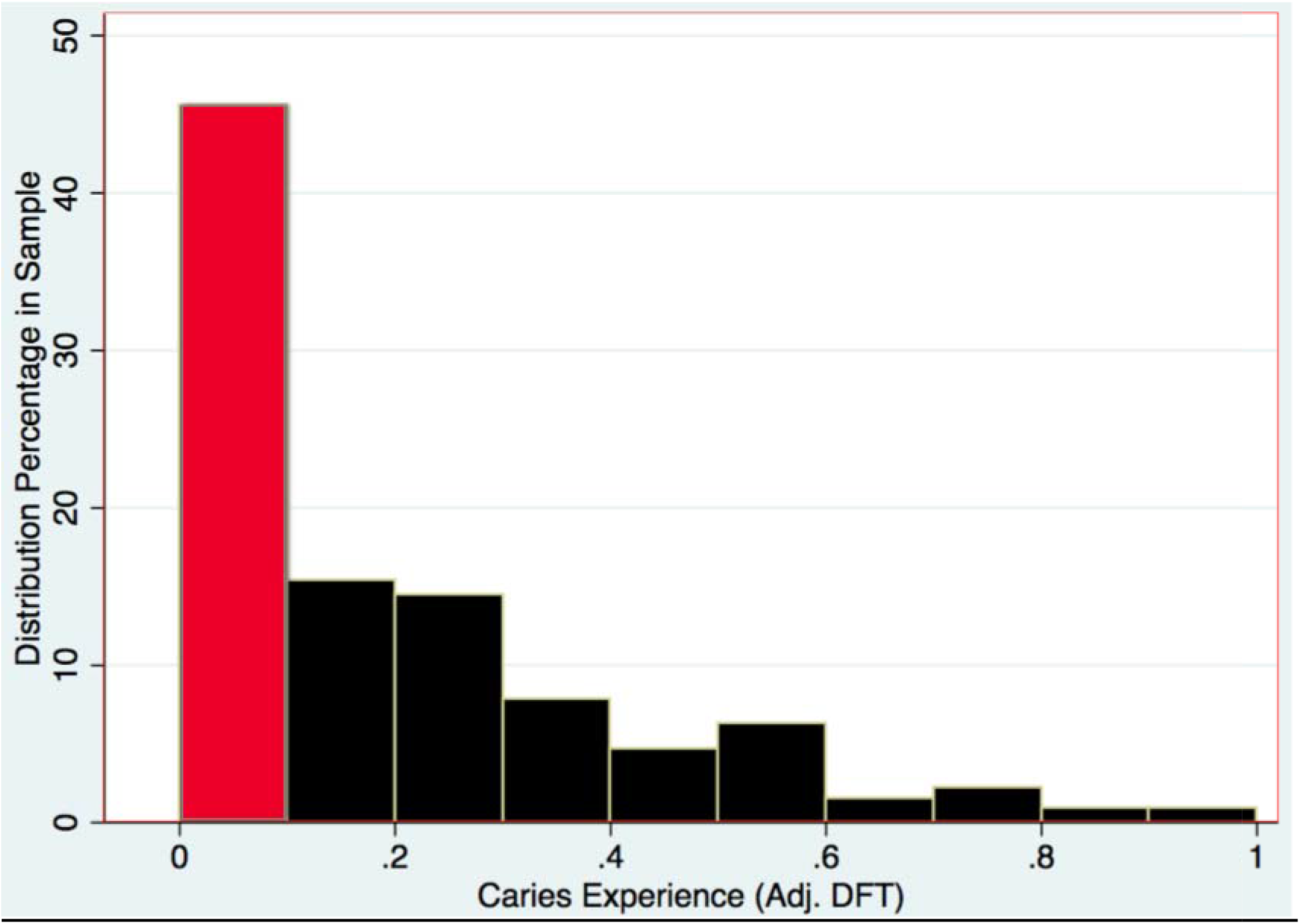
Distribution of continuous caries experience.

**Table 3.** Univariate logistic regression (AdjDFT=0 vs adjDFT>0)

|  | **n** | **OR (95%CI)** | **p** |
| --- | --- | --- | --- |
| Gender | 318 |  |  |
| Male |  | Ref | Ref |
| Female |  | 1.30 (0.80, 2.10) | 0.29 |
| Age (Years) | 318 | 1.09 (1.06, 1.14) | **<0.01** |
| OI type | 313 |  |  |
| I |  | Ref | Ref |
| III |  | 1.08 (0.57, 2.07) | 0.81 |
| IV |  | 0.84 (0.48, 1.45) | 0.53 |
| DI | 306 |  |  |
| No DI |  | Ref | **Ref** |
| DI Present |  | 1.93 (1.15, 3.25) | **0.01** |
| Debris index | 179 | 0.98 (0.58, 1.65) | 0.94 |
| Molar classification | 293 |  |  |
| I |  | Ref | Ref |
| II |  | 1.02 (0.47, 2.24) | 0.96 |
| III |  | 1.63 (0.94, 2.85) | 0.08 |

## Discussion

In this study we found that the prevalence of caries is in agreement with the findings of Malmgren^9^, as 30% of our OI population exhibits dental caries. O’Connell^8^ observed that while caries was rare in their study, some OI patients reported extensive dental restorations due to early childhood caries. In the present study, after incorporation of dental restorations and the total number of teeth present as a component, the caries experience was found to be present in 70% of the sample population (figure 2).

Individuals with primary, mixed and permanent dentition were represented in the study. Missing teeth were observed in a few cases, some being attributed to hypodontia or impacted teeth, while other cases presented with retained primary teeth. Use of the adjusted DFT to account for hypodontia enabled the consideration of retained teeth in the caries index, as well as facilitating the comparison within and across dentition-types (primary, mixed and permanent dentition). While the DT and FT scores of both OI types I and III are higher than type IV (table 1), an increase in CPE conforms to the order of increasing DI prevalence and severity in OI Types: Type I < Type IV < Type III (table 2). This difference in outcome may be attributed to the adjustment of the DFT score, which presents the dental-decay prevalence as a proportion of teeth affected by caries. This procedure overcomes the potential underestimation of the caries experience, by enabling more precise comparison of two groups with different number of teeth. Around 46% of the sample population (Figure 2) had negligible caries experience and caries distribution was skewed. The continuous variable CPE was converted into a binary variable (caries experience vs. no caries experience) and logistic regression was performed with the binary variable. This enables the result to approximate closely the outcome obtained if we could have utilized the DMFT index instead of adjusted DFT. In the OI population, a significant difference in CPE was found between groups with and without DI. In figure 3, the group with DI presents comparatively more CPE. Although caries experience is an irreversible disease, a dip in CPE is observed in the age group 8.5 to 13.7 years (figure 3). The age group coincides with the mixed dentition period and the dip is explained by the exfoliation of primary teeth and the eruption of newly erupted permanent teeth. Dental clinicians can be more inclined to provide preventive dental care such as pit & fissure sealants and fluoride application to OI individuals with DI especially in newly erupted permanent teeth to individuals in their mixed dentition age. The classification of OI type is done phenotypically, so the OI type may change over time as the manifestations change with age and bisphosphonate drug use. Because genetics is one of multiple of factors that contribute to the caries experience, future studies may require genotypic classification. Where different groups may exhibit different dental etiologies and possibly different caries experiences. In contrast to the present result of the study, dental caries may possibly be correlated with genetically classified OI.

**Figure 3.**
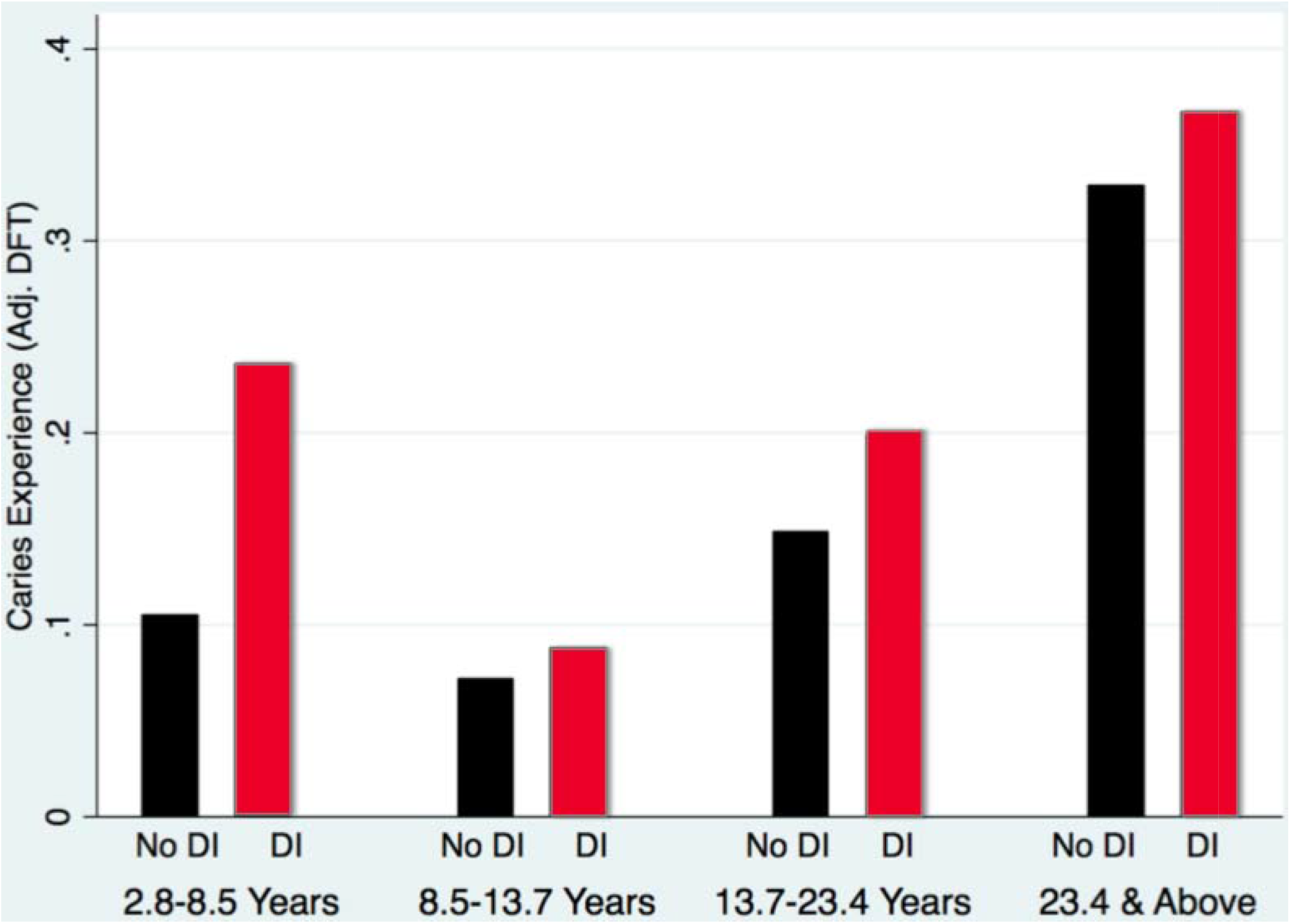
Adjusted DFT by DI status and age quartiles.

To our knowledge, this study is the first multi-center cross-sectional OI study to assess the caries experience of a large and diverse OI population. Within the limitations of this study, we provided preliminary data that identifies baseline characteristics to further formulate hypothesis for detailed analysis of a longitudinal data while controlling for socio-demographic and socio-economic predictors.

## Conclusion

We found no evidence that the CPE of OI patient is related to the OI types. Using an adjusted DFT, we found that the presence of DI increases the probability of developing caries experience when compared to subjects without DI while controlling for all other predictors of CPE.

